# Age-related Gene Expression Signatures (AGES) in rats demonstrate early, late, and linear transcriptional changes from multiple tissues

**DOI:** 10.1101/717835

**Authors:** Tea Shavlakadze, Melody Morris, Jian Fang, Sharon X. Wang, Weihua Zhou, Herman W. Tse, Ricardo Mondragon-Gonzalez, Guglielmo Roma, David J. Glass

## Abstract

In order to understand changes in gene expression that occur as a result of age, which might create a permissive or causal environment for age-related diseases, we produced a multi-timepoint Age-related Gene Expression Signature (AGES) from liver, kidney, skeletal muscle and hippocampus of rats, comparing 6, 9, 12, 18, 21, 24 and 27-month old animals. We focused on genes that changed in one direction throughout the lifespan of the animal, either early in life (early logistic changes); at mid-age (mid-logistic); late in life (late-logistic); or linearly, throughout the lifespan. The pathways perturbed as a result of chronological age demonstrate organ-specific and more global effects of aging, and point to mechanisms that might be counter-regulated pharmacologically in order to treat age-associated diseases. A small number of genes were regulated by aging in the same manner in every tissue, suggesting they may be more universal markers of aging.

## Introduction

Aging is the strongest risk-factor for many serious diseases and co-morbidities, including cancer, heart disease, kidney disease, dementia, Alzheimer’s disease, frailty and sarcopenia (Armanios et al., 2015; Egerman and Glass, 2014; Johnson et al., 2013; Kirkland and Tchkonia, 2015; Niccoli and Partridge, 2012). Increasing evidence suggests that aging occurs in a regulated manner, and that perturbation of discrete cell signaling pathways can extend lifespan and delay age-related diseases and co-morbidities (Bitto et al., 2016; Chen et al., 2009; Flynn et al., 2013l; Harrison et al., 2009; Miller et al., 2011; Miller et al., 2014; Zhang et al., 2014).

Multiple analyses of age-related changes in gene expression have been conducted in various tissues in mice (Barns et al., 2014; Braun et al., 2016; White et al., 2015) and rats (de Magalhães et al., 2009; Ibebunjo et al., 2013; Shavlakadze et al., 2018). There have been other strategies for the development of molecular signatures of aging; for example, researchers have determined DNA methylation patterns that correlate with biological age. An advantage of this method is that easily accessible tissue such as blood can be used to determine and cross-compare a human’s aging status (Gross et al., 2016; Sziráki et al., 2018; Thompson et al., 2018).

It is possible to obtain a more thorough molecular profile of aging by examining gene expression changes in multiple tissues throughout multiple time points in the lifespan of the animal, and determining genes that are perturbed in a consistent direction - in other words, genes that consistently increase in expression throughout life, or genes that consistently decrease in expression.

We were curious to know whether examination of multiple tissues at multiple time points could lead to new insights into the possible global and tissue-specific mechanisms of aging which might be causal for age-related pathologies. Thus, we undertook a study of gene expression changes with age throughout the animal’s lifespan (at 6, 12, 15, 18, 21, 24 and 27 months) in liver, gastrocnemius muscle, kidney and hippocampus. Rats were chosen because we had previously shown that rats are an excellent model for sarcopenia - the age-related loss of skeletal muscle (Ibebunjo et al., 2013). Here we are applying that experience to other tissues to develop a full multi-tissue aging signature. We discovered genes that change in common in every tissue; genes which are regulated in early-logistic, mid-logistic, late-logistic and linear fashion in particular tissues, and pathways that are regulated in multiple tissues, giving some indication of common mechanisms of aging. It is our hope that this dataset will serve as a valuable resource for further molecular insights into mechanisms of aging.

## RESULTS

### Transcriptional profiling of liver, gastrocnemius muscle, kidney, and hippocampus throughout the rat lifespan

We sought to establish both tissue-specific and more global aging gene signatures, which could serve as a basis for understanding the overall aging process. We were interested in genes that changed in a particular direction (either consistently up-regulated or consistently down-regulated) throughout the lifespan of the animal. Sprague-Dawley (SD) rats live approximately 2.5 – 3 years under laboratory conditions (Sengupta, 2013). The mortality rate of these rat cohorts at 21 months is ∼22%, at 24 months is ∼30% and at 27 months **∼**50% (Shavlakadze et al., 2018). In order to identify genes whose expression pattern is influenced by age, we performed gene expression profiling by RNAseq of liver, gastrocnemius muscle, kidney and hippocampus from male SD rats aged 6, 9, 12, 18, 21, 24 and 27 months. The gene expression data are available at the NIH Sequence Read Archive under the BioProject accession number PRJNA516151, and **Supplementary table 1A-D** contains the Log2 fold change and respective p values (raw and adjusted for the false discovery rate, FDR) for all genes in searchable excel format. Genomic data are visually presented as Volcano plots in **Supplementary figure 1**. PCA plots are shown in **Supplementary figure 2**, demonstrating that age distinguishes the genes from the various tissues, especially at the later time points. As a validation of the RNAseq analysis, we confirmed the reproducibility of age-related changes in genes that we have previously compared between young (6 month) and old (24 month) kidneys by RT-qPCR (Shavlakadze et al., 2018). 26 out of 28 previously validated genes reproduced in the present study (Shavlakadze et al., 2018). These include genes from the interferon gamma pathway (Cdkn1a, Psmb8, Vcam1, Cd38, Il7, Lgals3bp, Ube2l6, St8sia4, Il2rb, Isg20, Lcp2, Samhd1, C1s, Oas2, Ifitm3, Stat4) and genes from the “epithelial to mesenchymal transition” pathway (Col1a1, Col1a2, Col3a1, Col4a1, Spp1, Cd44, Fn1, Pdlim4, Timp1, Gja1).

### Age-regulated genes that followed a linear pattern, or early-, mid- or late-logistic patterns and gene pathways enriched by these genes

One of the key strengths of our dataset is the multiple age time-course obtained across the lifespan of the rats from four tissues. This dataset allowed us to identify age-regulated genes and their behavior patterns, in a manner distinct from multiple prior studies, where analysis was confined to “young” vs “old” samples. Age-regulated genes were screened for according to the following criteria (**Figure 1**, left side of the flow chart): a fold change of >1.5 (in either direction); Benjamini-Hochberg adjusted p-value < 0.05 for expression regulation between 6 month and at least one older age; and a Bayesian Information Criterion (as specified in the Methods), for the linear or four-parameter logistic fit that was at least 5% improved over that of the null model. Tracking expression levels of these genes throughout the rat lifespan allowed us to remove genes whose behavior was not consistent over time.

**FIGURE 1.**
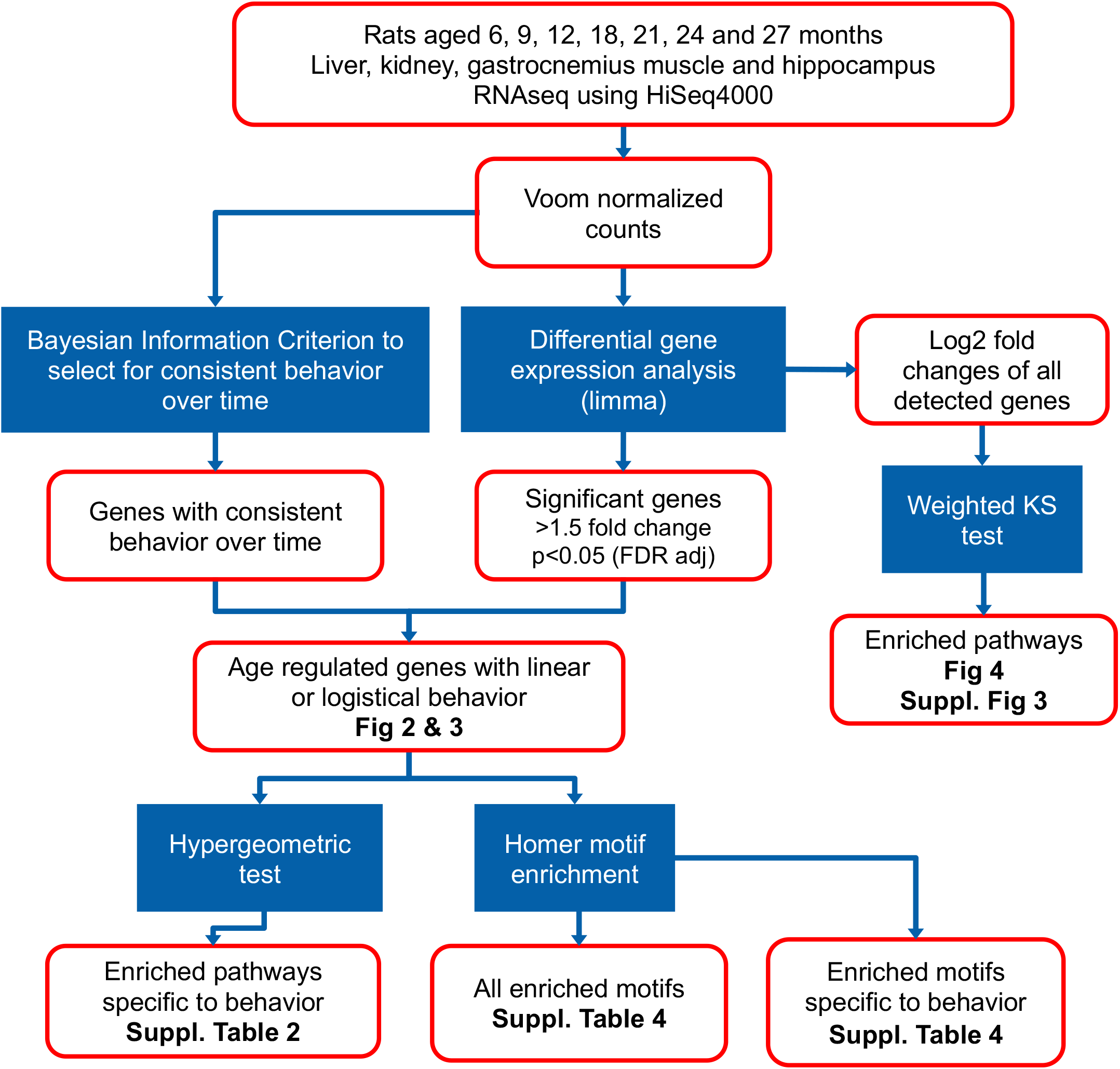
Flowchart for analyses used in the study. RNAseq data were generated using HiSeq4000 platform from liver, kidney, skeletal muscle and hippocampus of rats, comparing 6, 9, 12, 18, 21, 24 and 27-month old animals. The resulting counts were first normalized with the voom method and differential expression determined with limma. Differentially expressed genes were defined by the following criteria: a fold change of >1.5 (in either direction) and Benjamini-Hochberg adjusted p-value < 0.05 between 6 month and at least one older age. In addition to these criteria, the Bayesian Information Criterion of the linear or four-parameter logistic fit was compared to the null model (no relationship over time) to ultimately determine age-regulated genes. The age-regulated gene lists were used to perform pathway enrichment analyses that follow a linear pattern, or early-, mid- or late-logistic patterns and transcription factor motif enrichment (left side of the flowchart). For an alternative analyses arm, log2 fold changes from all detected genes were used for pathway enrichment by applying a weighted Kolmogorov–Smirnov (KS) test (right side of the flowchart).

In order to define the behavior of age-regulated genes, we chose a strategy that would allow us to catch genes that changed over a discrete interval, and then maintained their new set-point from there on. The behavior of the age-regulated genes was best fit by a linear or four parameter logistic model - again, to determine genes which changed in either a linear pattern, or early-, mid- and late-logistic patterns, where the point of inflection for the regulation in expression coincided with early (< 12 months), mid (≥12 months and <21 months) or late (≥21 months or older) life, respectively (see **Figure 3B-E** for examples). Though this analysis, we identified 1291 age-regulated genes in the liver, 1854 genes in the gastrocnemius muscle, 2195 genes in the kidney, and 229 genes in the hippocampus whose expression changed in a particular direction (either upward or downward) throughout the animals’ lifespan (**Figure 2 A, B, C, D**).

**FIGURE 2.**
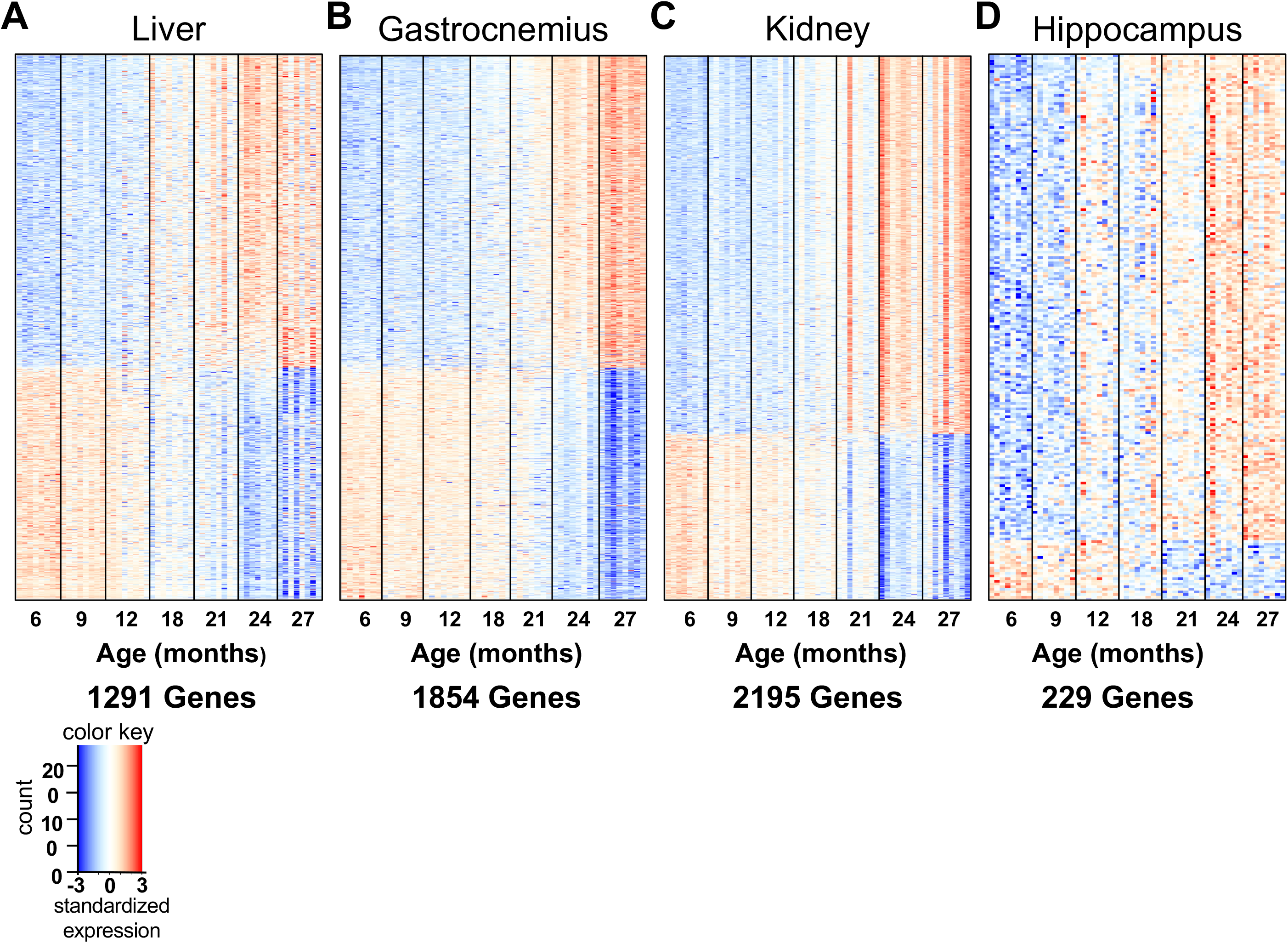
Heat maps depicting expression of age-regulated genes in liver (A), gastrocnemius muscle (B), kidney (C) and hippocampus (D) from rats aged 6, 9, 12, 18, 21, 24 and 27 months. There were 1291 age-regulated genes in the liver (A); 1854 in the gastrocnemius muscle (B); 2195 in the kidney (C); and 229 in the hippocampus (D). In heat maps, each row represents a single gene and each column represents a single rat (n=7-9 rats per group). For each gene (in each tissue), standardized expression levels were calculated by subtracting the mean expression level from each individual measurement and dividing by the standard deviation across all samples. The color key shows that the scaling of the standardized expression from low to high are colored from blue to red. Thus, gene up-regulation is shown by a transition from blue to red, and down-regulation by a transition from red to blue.

**FIGURE 3.**
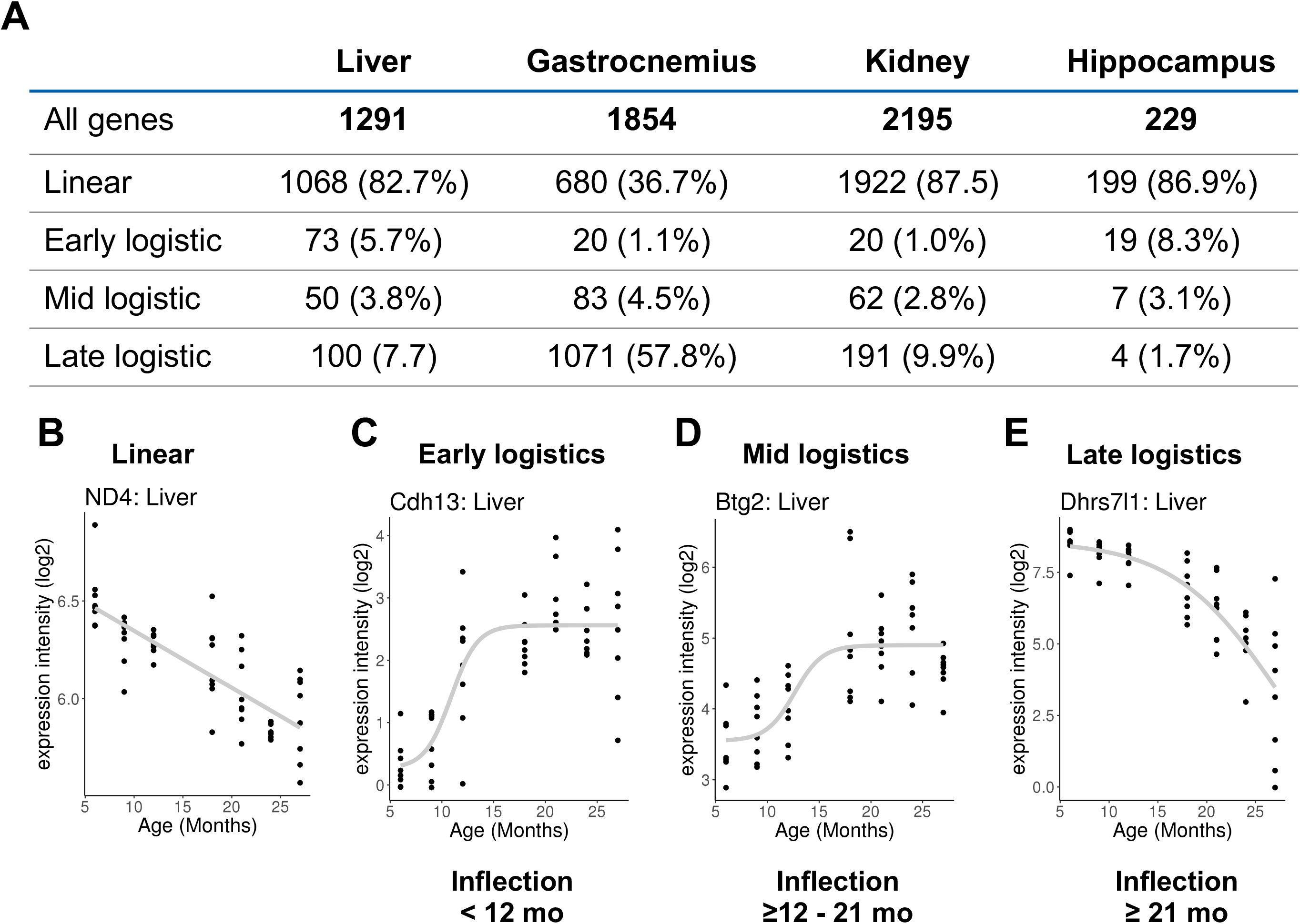
Numbers and percentages of age-regulated genes that followed a linear pattern, or early-, mid- or late-logistic patterns (**A**). Age-regulated genes were screened for according to the following criteria (**Figure 1**, left side of the flow chart): a fold change of >1.5 (in either direction); Benjamini-Hochberg adjusted p-value < 0.05 for expression regulation between 6 month and at least one older age; and a Bayesian Information Criterion (as specified in the Methods), for the linear or four-parameter logistic fit that was at least 5% improved over that of the null model. These genes were identified based on data sets from liver, gastrocnemius muscle, kidney and hippocampus across 7 ages: 6, 9, 12, 18, 21, 24 and 27 months, n=7-9 rats per group. Example genes demonstrating a linear (**B**), early logistic (**C**), mid-logistic (**D**), or late-logistic (**E**) patterns are included.

When analyzed by logistic or linear behavior in the liver, kidney, and hippocampus, the majority of the age-regulated genes changed in a linear fashion: 82.7%, 87.5% and 86.9% of all age-regulated genes respectively (**Figure 3A**). In contrast, the majority of age-regulated genes for the gastrocnemius muscle followed a late-logistic pattern (57.8%), with 36.7% of the genes being linear (**Figure 3A**).

We next sought to identify pathways that were enriched by genes with the linear, early-, mid- and late-logistics behavior patterns. This was achieved by performing hypergeometric enrichment test on genes with the linear, early-, mid- and late-logistic behavior against the background of all genes that changed with age in at least one tissue (**Supplementary table 2**).

This method would identify pathways that stand out above the background of the global age-related transcriptional regulation. According to the behavior type, the majority of the identified pathways were “linear up-regulated” pathways and these were seen in the kidney, liver and hippocampus. Among the “linear up-regulated” pathways, those related to the immune response were shared between kidney and hippocampus (**Supplementary table 2**). Pathways related to cholesterol and lipid metabolism increased linearly in the aging liver (**Supplementary table 2**).

Early response pathways were enriched in the gastrocnemius muscle and were predominately driven by transcription factors Fos, Jun, Erg1 (**Supplementary table 2**). These early response genes are upregulated in middle aged rats (by 12 months of age), which corresponds to ∼45 years in humans and it is possible that these factors are responsible for initiating later events.

Interestingly, transcription factors Fos, Jun and Erg1 are activated downstream of Ras/Raf/MEK/ERK signaling and pharmacological inhibition of this pathway has been shown to extend lifespan in *Drosophila* (Slack et al., 2015). In the gastrocnemius muscle, we also found up-regulated (myogenesis) and down-regulated (glucose and carbohydrate metabolism) pathways, which changed with a late logistic profile (**Supplementary table 2**). Increase in the myogenesis pathway in late life is likely to be related to functional denervation of old myofibers, as well as regenerative events in response to age-related myofiber deterioration.

### Biological pathway enrichment of gene expression changes with age in liver, gastrocnemius muscle, kidney and hippocampus

In addition to the pathway enrichment based on age-regulated genes described above, we also performed a more global pathway enrichment analyses comparing 6 month old rats to older ages (i.e. 9 *vs* 6 months; 12 *vs* 6 months, etc. and ultimately 27 *vs* 6 months) (**Figure 1**, right side of flow chart). Mean fold-change and statistical significance of genes in enriched pathways are shown in **Figure 4A and 4B**, respectively. A complete list of enriched pathways based on statistical significance is provided in **Supplementary figure 3.** We observed that many of these pathways were common to at least three tissues. Furthermore, while the fold change of gene expression enriched to each pathway was lower in hippocampus compared with liver, gastrocnemius and kidney (**Figure 4A**), the statistical significance of the skew of the pathways towards up or down regulation was similar for several pathways (**Figure 4B**). Most strikingly, pathways which were related to upregulation of inflammation were the dominant theme that we observed; for example, pathways related to the innate immune response, inflammation and cytokine signaling were up-regulated with age in the liver, gastrocnemius and in the kidney (**Figure 4A and 4B**). Age-related upregulation of these pathways was also seen in hippocampus, albeit with a less dramatic increase. Pathways linked to Allograft Rejection and Interferon Alpha and Gamma Response increased strongly with age in kidney, liver and gastrocnemius muscle, and were also increased in hippocampus (**Figure 4A** and **4B**). The Complement pathway was also up-regulated with age in all four tissues. In addition to inflammation, other pathways of interest were perturbed; the Apoptosis pathway was overall up-regulated in aging liver, gastrocnemius and kidney and to a lesser extent in hippocampus, suggesting an overall increase in cell death occurring in tissues with age - perhaps this could be a consequence of loss of growth factors, or other positive anabolic signals. Metabolic pathways were generally downregulated with age in the liver, skeletal muscle and kidney, except for the cholesterol homeostasis pathway, which gradually increased in the liver with aging (**Figure 4A and 4B**).

**FIGURE 4.**
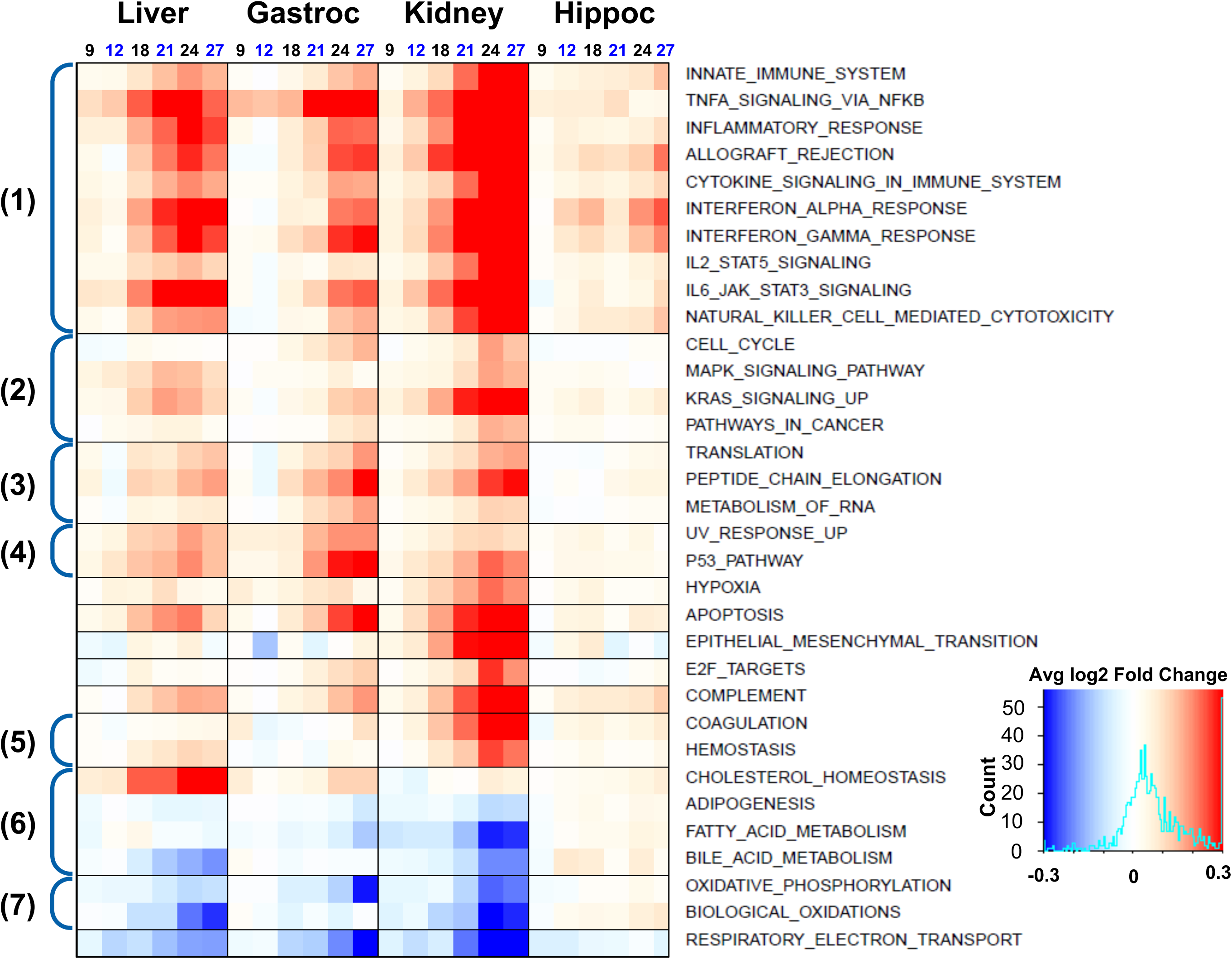
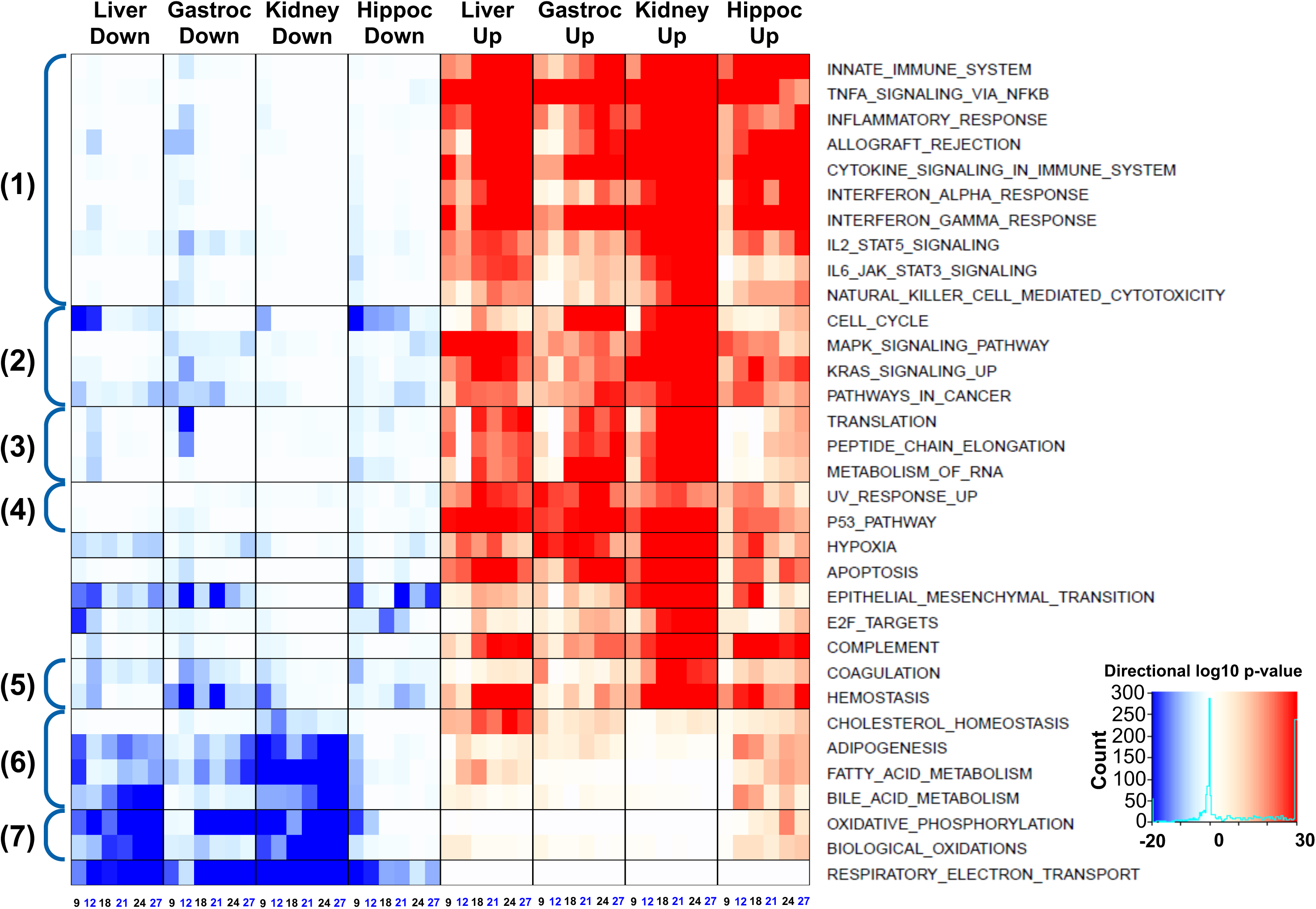
Identification of pathways UP- or DOWN- regulated by aging. Biological pathway enrichment for gene expression changes with age in liver, gastrocnemius muscle, kidney and hippocampus. In both A and B, each row represents a pathway, annotated on the right hand side and each column represents a specific age comparison (i.e. 9 months vs 6 months, 12 months vs 6 months, etc up to 27 months vs 6 months). Pathway analyses are based on n=7-9 rats per group. Colors represent average log2 of the fold change of genes in the indicated pathway (A) or statistical significance of enrichment of the pathway (B). Color transition from white to red shows up-regulation, and color transition from white to blue shows down-regulation of the pathway. The color key in (A) indicates the corresponding average log2 of the fold change and the color key in (B) indicates statistical significance of log10 of the p-value of enrichment. Numeric annotation on the left hand side (A,B) indicates grouping of pathways by themes: (1) Inflammatory; (2) Growth; (3) RNA Processing; (4) DNA Damage; (5) Coagulation; (6) Metabolic; (7) Biological oxidation.

The most prominent pathways down-regulated with aging were mitochondrial: oxidative phosphorylation, respiratory electron transport and biological oxidation were all gradually down-regulated with age in the liver and kidney. Prominent downregulation of oxidative phosphorylation and respiratory electron transport was seen in the gastrocnemius muscle. Age-related regulation of oxidative phosphorylation, respiratory electron transport and biological oxidation pathways was not conspicuous in hippocampus (**Figure 4A** and **4B**). These changes were consistent with the idea that mitochondria become less competent with age, depriving cells of critical supplies of ATP, in addition to the multitude other signals, which are mitochondrial in origin.

### Common age-regulated genes among multiple tissues

We also sought to determine whether there were common age-regulated genes among the multiple tissues and identified 148 genes that were common to at least three of these tissues (here referred to as the “common genes” **Supplementary table 3**). The majority of common genes (n=110) were upregulated with aging consistently across tissues. Twenty-five common genes had a mixed expression pattern, i.e. were up-regulated in some tissues and down-regulated in others. Thirteen common genes were down-regulated in at least three tissues (liver, gastrocnemius and kidney). Interestingly, expression of 13 common genes was up-regulated with age in all four tissues, with 11 of these genes being annotated: LOC103689965, Psmb8, Gpnmb, Tspo, Irf7, Icam1, Ms4a6a, Isg15, C4a, C4b, Cebpb (**Table 1**, and the first 13 transcripts in **Supplementary table 3**).

**TABLE 1.** Age-regulated genes that were in common to 4 tissues and were up-regulated with age in a linear manner. For each gene, the most significant adjusted p-value differentiating 6 month from older ages is shown. A complete list of age-regulated genes that were in common to at least 3 tissues is provided in **Supplementary table 3**.

|  | Behavior |  |  |  | The most significant adjusted p-value differentiating 6mo from later ages |  |  |  |
| --- | --- | --- | --- | --- | --- | --- | --- | --- |
| Symbol | Liver | Gastroc | Kidney | Hippoc | Liver | Gastroc | Kidney | Hippoc |
| LOC103689965 | linear | linear | linear | linear | p<2.33E-07(24Mo) | p<5.12E-06(24Mo) | p<1.26E-08(27Mo) | p<8.52E-06(24Mo) |
| Psmb8 | linear | linear | linear | linear | p<1.98E-04(24Mo) | p<1.45E-04(24Mo) | p<1.97E-07(27Mo) | p<4.09E-03(24Mo) |
| Gpnmb | linear | mid logistic | linear | linear | p<1.87E-06(27Mo) | p<6.24E-15(24Mo) | p<9.94E-07(27Mo) | p<7.32E-08(24Mo) |
| Tspo | linear | late logistic | linear | early logistic | p<7.52E-04(27Mo) | p<2.11E-13(24Mo) | p<7.22E-09(27Mo) | p<1.58E-03(27Mo) |
| Irf7 | linear | linear | linear | linear | p<1.14E-04(27Mo) | p<2.50E-02(24Mo) | p<2.31E-04(27Mo) | p<2.39E-03(24Mo) |
| Icam1 | linear | linear | linear | linear | p<1.25E-07(27Mo) | p<2.27E-03(24Mo) | p<2.23E-08(27Mo) | p<1.41E-03(27Mo) |
| Ms4a6a | linear | linear | linear | linear | p<1.81E-03(24Mo) | p<9.21E-04(24Mo) | p<5.04E-08(27Mo) | p<3.17E-02(24Mo) |
| Isg15 | linear | linear | linear | linear | p<1.28E-03(27Mo) | p<1.58E-02(24Mo) | p<1.25E-03(27Mo) | p<2.25E-02(24Mo) |
| C4b | linear | linear | linear | linear | p<1.62E-07(24Mo) | p<1.49E-04(24Mo) | p<2.13E-08(27Mo) | p<4.00E-04(24Mo) |
| C4a | linear | linear | linear | linear | p<2.20E-07(24Mo) | p<3.86E-06(24Mo) | p<1.37E-08(27Mo) | p<7.64E-06(24Mo) |
| Cebpb | linear | linear | linear | linear | p<2.41E-03(24Mo) | p<1.49E-07(24Mo) | p<1.57E-04(24Mo) | p<1.07E-02(27Mo) |
| ENSRNOG00000059770 | linear | late logistic | linear | linear | p<2.52E-02(27Mo) | p<1.60E-08(24Mo) | p<3.87E-04(27Mo) | p<4.46E-03(27Mo) |
| ENSRNOG00000059927 | linear | linear | linear | linear | p<2.17E-03(24Mo) | p<4.69E-07(24Mo) | p<1.96E-07(27Mo) | p<1.42E-02(24Mo) |

In analyzing the gene changes over time, we were surprised to see another characteristic emerge from the data - interestingly, aging induced an increase in the global gene expression variability (**Supplementary figure 4**). Density plots of standard deviation variability demonstrated higher overall gene expression variability in 27-month old liver, gastrocnemius and kidney, but not hippocampus, compared with samples from 6-month old animals (**Supplementary figure 4**). No increase in gene expression variability in the old hippocampus is in line with an overall more muted response to aging seen in this tissue in aged Sprague Dawley rats (**Figure 2D**).

### Transcription factor motif enrichment in liver, gastrocnemius muscle, kidney and hippocampus

Given the sets of genes which were observed to be regulated by age, the next question was, what upstream mechanisms might be perturbing these genes? We therefore next sought to assemble the set of transcription factors that might be responsible for the gene changes, by using an program to identify transcriptional binding site motif enrichment, known as HOMER (for Hypergeometric Optimization of Motif EnRichment) (Heinz et al., 2010). HOMER can be used to identify motifs that are statistically enriched in the promotor region of a given list of genes. We first applied HOMER using motifs of 6-12bp length over a range of 1000bp upstream and 50bp downstream of each transcriptional start site, using the age-regulated gene list for each tissue (**Figure 1**, left side of the flowchart). **Supplementary table 4A** contains the list of these enriched motifs (p-value<0.05 adjusted by Benjamini-Hochberg Procedure). In addition, we applied the HOMER program to the lists of gene subsets with linear, early-, mid- and late-logistic behaviors, here defined as age-regulated genes (**Supplementary table 4B).**

Overall, the most prominent binding motifs enriched by genes up-regulated with age were transcription factors linked with the innate immune response and inflammation. For example, the motif for the transcription factor IRF2 (interferon regulatory factor) was enriched by promoters of up-regulated genes in three tissues, liver, kidney and hippocampus, consistent with the finding that the interferon signaling pathway was found to be up-regulated in all four tissues coincident with age - but this motif commonality allows focus on IRF2 in particular (**Supplementary table 4A)**. Genes that contributed to the identification of the IRF2 motif were characterized by linear up-regulation with aging (**Supplementary table 4B**).

The motifs associated with IRF1, IRF3, and ISRE (interferon stimulated response element) were enriched by gene promoters with linear behavior in the liver and kidney, where robust age-related induction of the interferon alpha and interferon gamma signaling occurred (**Supplementary table 4A,B**). The motifs associated with IRF8 were enriched in kidney and hippocampus (**Supplementary table 4A,B**).

Motifs related to the inflammatory response (STAT4, STAT5, STAT1 and NfkB) were also enriched by promoters of linearly up-regulated genes (**Supplementary table 4B**). Motifs related to STAT4, STAT5, STAT1 were prominent in the aging kidney and motifs related to NfkB were prominent in the aging liver (**Supplementary table 4B**). Furthermore, in the aging kidney, the motifs for hepatocyte nuclear factors (HNF4a, HNF1 and HNF1b) were enriched by promoters of down-regulated genes (**Supplementary table 4A, B**).

In the gastrocnemius muscle, the Mef2 a,b,c and d (myocyte enhancer factor-2) transcription factor related motifs were enriched by gene promoters that were down-regulated later in life coinciding with age-related muscle atrophy (**Supplementary table 4A and B**). Interestingly, in the gastrocnemius Mef2a related motif was also enriched by up-regulated gene promoters with early logistic behavior. Since early response genes assigned to this gene subset were differentially regulated between 9 and 12 months (middle age), up-regulation of factors that target the Mef2a motif may be linked with early events of muscle aging and may trigger later events of sarcopenia.

In gastrocnemius muscle, PRDM14 (PR domain zinc finger protein 1) and a zinc finger transcription factor Sp1 (specificity protein 1)-related motifs were enriched by promoters of up-regulated genes with a late-logistic behavior (**Supplementary table 4B**).

## DISCUSSION

There are multiple definitions of aging; our interest was to determine whether we could find a reproducible pattern of gene expression changes over the lifespan of a rat in multiple tissues to establish an Age-associated Gene Expression Signature of aging (denoted AGES). The power of profiling multiple ages, rather than only two (young versus old) was that it allowed for the demonstration of several distinct patterns of gene expression changes that may occur. Profiling several distinct organs at the same time allowed us to cross-compare the changes observed in each organ in order to learn which were unique and which were universal. We envision that AGES will help in research efforts to (1) identify the particular expression changes that are either diagnostic or causative of age-related disease; (2) identify changes that create an environment in which disease is more likely develop - for example by decreasing the efficiency of DNA damage repair, or by downregulating mitochondrial genes, or by upregulating genes whose protein products can induce inflammation-associated damage (3) identify early triggers of the aging process; (4) identify late-stage, disease associated biomarkers of aging and age-associated diseases. In addition, certain gene expression changes may be protective or compensatory, rather than deleterious for the aging process. And, of course, the hope is that AGES will identify pathways and mechanisms associated with disease that point to critical pathways and targets that could be considered for pharmacological intervention.

AGES should also help determine the relative efficacy and mechanisms by which true “anti-aging” drugs operate, by identifying that component of the signature which is counter-regulated by such a drug. For example, a drug like an antibiotic could be life-saving, but would not be expected to perturb AGES; however, a pharmacologic agent which extended lifespan in an “anti-aging” manner might be expected to counter-regulate critical components of AGES. This sort of result was seen when a rapalog, a class of drugs which has been shown to increase lifespan in many organisms(Harrison et al., 2009; Miller et al., 2011; Miller et al., 2014), was used in older rats, resulting in kidney protection and a reversal of a subset of age-regulated genes (Shavlakadze et al., 2018). Similarly, a six-month resistance training program in elderly humans that improved skeletal muscle strength was accompanied by a reversal of the transcriptional signature of mitochondrial function to a younger phenotype (Melov et al., 2007).

The time course used in this study demonstrated that gene expression changes fell into several distinct classes: “early logistic,” where the gene changed in expression relatively early in life; “mid logistic,” where the change happened in mid-life, and was then consistent; late-logistic, where the changes happened at a later time point, and may have been thus more coincidental with the onset of age-associated disease or pathology; and linear changes - these genes seemed to track chronological aging faithfully, and thus might be useful in understanding how particular genes “know” how old an animal is.

Among the pathways which were found to be linear in their behavior, most prominent were the various inflammatory pathways, including the increase in the complement pathway, and interferon signaling, most prominently in the kidney, but also in other tissues (**Supplementary table 2**). In the future. it will be interesting to see if kidney aging is a prominent driver in other organisms, as it seems to be in the Sprague Dawley rat. Given the prominence here, one is tempted to ask if kidney-specific perturbations would be sufficient to modulate aging in the entire organism. One hint of this was our recent study using a rapalog, which showed significant improvement in kidney pathology with short-term treatment in aged animals, coincident with counter-regulation of the aging pathways (Shavlakadze et al., 2018). Rapalog treatment has reproducibly induced an increase in lifepsan, even when given in older animals.

A follow-up question that these patterns raise is which set of genes might house the molecular signatures which induce the negative phenotypes associated with age. For example, the late-logistic genes are coincident with age-associated disease, such as sarcopenia (Ibebunjo et al., 2013); however, it may be that this set of gene changes is triggered by the genes that are perturbed earlier - for example in mid-logistic phase. Of course it is also possible that the genes which are regulated in a linear fashion throughout life may be the real “time-keepers”, and that there is an eventual threshold that is reached by these genetic regulations - a stage which triggers the negative sequalae of age. Determining the causative mechanisms triggering age-related gene expression will of course be the subject of many studies; it seems unlikely that there would be only a single inducing step, but it will still be useful to ask discrete questions, such as what is the mechanism for the induction of the dramatic increase in inflammatory gene expression seen in multiple tissues, and what is the mechanism for the decrease in mitochondria-associated gene expression?

As for the question, “does aging cause the same gene expression changes in all tissues?” there was considerable overlap in pathways perturbed by age in the kidney, liver and gastrocnemius muscle, but it was noteworthy to see many genes that were tissue-specific. This again brings up the challenge of conceiving of a single agent to broadly counter-regulate age-related gene changes, but it also highlights in particular gene pathways which are in common among the various tissues.

There were 13 genes regulated with age in all four tissues examined: LOC103689965, Psmb8, Gpnmb, Tspo, Irf7, Icam1, Ms4a6a, Isg15, C4b, C4a, Cebpb, a linc RNA previously unannotated and ENSRNOG00000059927, which apparently encodes for an unstudied protein whose sequence is consistent with it being an MHC1 receptor (**Table 1**). Several of these have been previouly shown to correlate with aging in specific tissues, however our data shed light on a potential systemic dysregulation of these genes and their associated functions. This is highly relevant as it provides insights for the potential mechanisms underlying the overall age-associated conditions. Consistent with the finding that the complement pathway as a whole is upregulated in aging, C4a, C4b and LOC103689965 (C4-like) are all components of this pathway. An enhanced activation of the classical and alternative complement pathways, as well as increased concentration of complement components in serum, has been observed in older (∼62 year) individuals compared to younger (∼26 years) ones. Although this correlation needs to be validated in a longitudinal study, other studies have shown elevated levels of complement components in aged brains of mice (Reichwald et al., 2009) and humans (Cribbs et al., 2012) - accumulation of complement member C1q occurs in areas of the brain vulnerable to neurodegeneration in aged mice, and knock out mouse models of the key components C1q and C3 display less age-related cognitive and memory decline (Shi et al., 2015; Stephan et al., 2013).

Age-related increase in Cebpb, Isg15, Icam1, Irf7, Psmb8 gene expression is likely to be linked with up-regulation of inflammatory pathways. In the brain, these changes are suggestive of age-related induction of neuro-inflammation and activation of glia. C/EBPβ (coded by the Cebpb gene) is a basic-leucine zipper DNA-binding protein that controls expression of cytokines and other pro-inflammatory genes, and induces pathways that are critical for glia activation (Bradley et al., 2003; Wang et al., 2018). Apart from being implicated in neuro-inflammation, C/EBPβ controls expression of delta-secretase enzyme, which is thought to contribute to the pathology in Alzheimer’s disease by cleaving both Tau and beta-amyloid precursor protein (Wang et al., 2018). ISG15 (interferon stimulated gene 15) (Kessler et al., 1988), which also was upregulated in every tissue examined, is a ubiquitin-like protein, and its expression is induced by type-I IFNs and by p53 (Huang and Bulavin, 2014) (Park et al., 2016). Under DNA damage conditions, ISG15 conjugates to p53 (referred to as p53 ISGylation): this modification enhances the binding of p53 to the promoters of its target genes, perhaps leading to suppression of cell growth and tumorigenesis. Although in aging increased binding of p53 to DNA can induce a senescent phenotype (Hafner et al., 2019).

While the data from liver, kidney and gastrocnemius muscle indicated that many genes were regulated with age, we identified far fewer age-regulated genes in hippocampus (229 genes). Furthermore, the response to age was muted in the hippocampus compared to the other tissues in that both the magnitude of fold changes and the statistical significance were smaller in hippocampus compared with other tissues (**Supplementary Figures 1)**. We have also observed this muted response in an earlier study that compared an independent cohort of 24-month old and 6-month old rats (data not shown). To interrogate the RNAseq results from hippocampus, we performed a side-by-side analyses of this earlier study and the current time-course study and identified genes that were differentially regulated with a fold change of > 1.5 (Benjamini-Hochberg adjusted p<0.05) between 6 and 24 months. We found 121 differentially regulated genes (108 were up-regulated and 13 were down-regulated) in the earlier study and 226 differentially regulated genes (210 were up-regulated and 16 were down-regulated) in the current study and there were 53 up-regulated genes (p<1e-70) and 7 down-regulated genes (p<1e-18) in common to the two studies. These results indicate that the smaller age-associated gene expression signature in the hippocampus is driven by the biology of the tissue, at least in the Sprague Dawley rat.

An interesting observation which was especially evident in the latest time points was that aging induces an increase in variability of gene expression. At the 27 month time point, two different classes of animals became evident - one which was “genetically younger” than the other, since its gene expression pattern maintained a greater consistency with earlier time points, and one which was “genetically older”, since its expression pattern continued along the age-associated slope established by the earlier time points. Other studies have pointed to this increased variability as well (Barns et al., 2014). Potential causes include an incursion of new tissue types in organs where they are not usually extant - for example age-associated fibrosis is a common event, and fibroblasts would be expected to derail an organ’s gene expression signature if these cells are not normally a significant component of that tissue type. Dysregulation of heterochromatin, allowing for expression from previously-silent chromosomal regions, could also contribute to this variability (Fushan et al., 2015).

Since aging is the biggest risk factor for the most serious diseases that affect humans, including cancer, heart disease, dementia, sarcopenia and frailty, and chronic kidney disease, it seems plausible that the gene-changes associated with aging could contribute to the onset of these conditions, either by creating a permissive environment for further pathology, or by directing causing aspects of these diseases. Pathways which were in common to all four tissues include the inflammatory pathways, especially interferon alpha and interferon gamma responsive genes, JAK-STAT activated genes, and TNF-alpha induced genes; these were up-regulated in all tissues- and most strongly in the kidney. From the clinical perspective, it is noteworthy that up-regulation of gene pathways induced by inflammatory cytokines and interferon gamma is also a characteristic to the human kidney aging (O’Brown et al., 2015).

Pathways which were downregulated in liver, gastrocnemius muscle and kidney included mitochondrial-pathways: oxidative phosphorylation and respiratory electron transport pathways. This highlights again the point that aging in part may be due to a loss of mitochondrial competence, a point which has been made many times in the literature. Here we show that a dramatic decline in mitochondrial competence associated pathways occurs later in life, meaning that one might hope that intervention rather late in life could still be quite helpful, before the onset in the dramatic downturn in mitochondrial competence.

A search for transcription factor motifs in the promoter regions of the genes which were regulated by aging causes further focus on interferon signaling (**Supplementary table 4**). Enhanced use of the IRF2 motif was found to be significantly associated with three tissues, kidney, liver and hippocampus. Several other Interferon Response factor motifs - IRF3, IRF8, IRF1 and ISRE - were associated with aging in liver and kidney. This repeated finding of interferon gamma and alpha signaling increasing in age, in multiple tissues, using multiple modes of analysis, does cause particular focus on these pathways - leading one to ask whether discrete inhibition of interferon signaling in aged animals might benefit age-related disorders. The finding also leads one to ask for the mechanism causing this reproducible increase. One idea is that the increase in LINE-1 element transcription, recently shown (De Cecco et al., 2019), could cause double-stranded RNA production, giving cells the impression they are being infected by viral agents.

We hope to continue to build on AGES by profiling more organs and cell-types, continuously building on the current AGES. Of course, this will be a long and continuous task, given the multiple additional organs that could be eventually added to the signature, including heart, lungs, hematopoietic and lymphoid cells, blood vessels, sensory organs, etc. Also, with the advent of single-cell RNA profiling, and given a desire for increasing granularity to help tease out the mechanisms which might bridge gene changes between early- mid- and late logistic genes, the idea of a truly complete AGES seems daunting indeed. For now, the current version should provide a rich resource to inspire new questions for further follow-up and study.

## METHODS

### Animal maintenance and tissue collection

All procedures involving animals were approved by the Institutional Animal Care and Use Committee of the Novartis Institutes for Biomedical Research (NIBR), Cambridge, MA, USA. Male Sprague Dawley (SD) rats were purchased at 3-4 weeks of age from Envigo (Indianapolis, USA) and aged at Envigo under specific pathogen free (SPF) conditions. Rats used for this time-course study were from cohorts with different birth dates, however rats in the same age group had the same week of birth. Rats were imported to NIBR animal holding facilities 4-8 weeks prior to tissue collection. Once imported, rats were housed singly and maintained at the SPF facility with controlled temperature and light (22°C, 12-h light/12-h dark cycle: lights on at 0600h/lights off at 1800h) and with ad libitum access to food (2014 Teklad Global 14% Protein diet, Envigo) and water *ad libitum*. Tissue collection occurred over several months and tissues form the same age group (n=8-10 per group) were collected on the same day between 12pm and 2pm to control for circadian variations. Prior to tissue collection, rats were fasted for 6 hours (with ad libitum access to water), from 0600 h to 1200 h, anesthetized with 3.5% isoflurane and killed by exsanguination and thoracotomy. Tissues were collected and frozen in liquid nitrogen.

### RNA extraction and integrity assessment

Snap frozen livers, gastrocnemius muscles, kidneys and hippocampi were ground in liquid nitrogen by mortar and pestle and total RNA was extracted from ∼30 mg of tissue powder using miRNeasy Micro Kit (Qiagen, 217084). The RNA concentration was quantified using NanoDrop Spectrophotometer (NanoDrop Technologies, USA) and the integrity assessed by the OD260/OD280 absorption ratio (>1.8) and by RIN score (≥8) using Agilent 2100 Bioanalyzer, RNA 6000 Nano LabChip kit and Agilent 2100 Expert Software (Agilent Technologies, Inc., Santa Clara, CA, USA). Samples with RIN scores ≥8 were designated for RNAseq.

### Transcriptomic analyses with RNAseq

Gene expression levels were measured using RNA sequencing technology. The RNA libraries were prepared using the Illumina TruSeq Stranded Total RNA Sample Preparation protocol with the Ribo-Zero Gold Kit and sequenced using the HiSeq4000 platforms following the manufacturers protocol. Samples were sequenced in paired-end mode to a length of 2×76bp base-pairs. Images from the instrument were processed using the manufacturers software to generate FASTQ sequence files. Read quality was assessed by running FastQC on the FASTQ files. Sequencing reads showed excellent quality, with a mean Phred score higher than 30 for all base positions.

For read alignment to the rat (*Rattus norvegicus*) genome and gene expression quantification for genes annotated by Ensembl v6.0.87^1^, the exon quantification pipeline was used (Schuierer and Roma, 2016). **Supplementary table 5** summarizes reads sequenced and mapped for each tissue. For each tissue, any gene with an average voom-normalized expression value of less than zero in any age group was considered undetected and removed from further analysis. As part of the initial QC, we performed a principle components analysis on the gene expression data from each tissue. However, we noticed that, in few cases, a single sample contributed overwhelmingly to the variability of any of the first 5 PCs. These samples were considered technical outliers and were removed from further analysis. **Supplementary figure 2** shows principal component analyses that best correlated with age for liver, gastrocnemius muscle, kidney and hippocampus datasets.

### Definition and characterization of age-regulated genes

Differential expression analysis was performed on the counts using voom normalization followed by limma (Ritchie et al., 2015) for model-fitting and contrast calculation with R version 3.4.2 (**Figure 1**). Because we had profiled multiple ages in the animals lifespan, we sought to characterize gene behavior as following a linear or logistic trend. To assess the overall gene behavior, we used Bayesian Information Criteria (BIC) to determine if the expression of a gene as a function of time was best fit by a linear model, a four-parameter logistic model, or a “null” model (no change over time).

Age-regulated genes were defined as those that met the following criteria: a fold change of >1.5 (in either direction) and Benjamini-Hochberg adjusted p-value < 0.05 for expression regulation between 6 month and at least one older age in addition to a BIC for the linear or four-parameter logistic fit that was at least 5% improved over that of the null model.

### Pathway enrichment analysis for age-regulated genes that specifically follow linear pattern, or early-, mid- or late-logistic patterns with aging

For each tissue, pathway enrichment analysis was performed on subsets of age-regulated genes that followed linear, early-, mid- or late-logistic patterns (**Figure 1**, left side of the flowchart). For each canonical pathway or hallmark pathway, the corresponding gene set was compared with an up- or down-regulated behavioral gene list. The overlapping genes were counted, and a hypergeometric test was performed to measure the statistical significance, e.g. p-value, on whether the number of overlaps occurred by chance when comparing the background of all genes included in the study. Finally, adjusted p-values were derived by the Benjamini-Hochberg procedure to control the FDR. We further repeated the same hypergeometric test on each behavioral gene list but using the background of all genes that changed with age in at least one tissue. The pathways enriched by both backgrounds were identified to be specific to linear, early-, mid- or late-response among the age-regulated genes.

### Biological pathway enrichment analyses of gene expression changes with age

For this analysis, up- or down- regulated biological pathways for each time-point comparison were assessed (**Figure 1**, right side of the flowchart). Canonical pathways (c2.cp.v5.2.entrez.gmt) and hallmark pathways (h.all.v5.2) were downloaded from the canonical signatures database and the hallmark signatures database (http://software.broadinstitute.org/gsea/msigdb/). For each tissue, the time-point comparisons considered were 9- vs 6-month, 12- vs 6-month, 18- vs 6-month, 21- vs 6-month, 24- vs 6-month, and 27- vs 6-month. For each pathway and time-point comparison, a weighted Kolmogorov–Smirnov (KS) test was performed as previously described (Kristinsson et al., 2012). The pathways (**Supplementary figure 3** and **Figure 4A,B)** were selected based on the following criteria: the p-value of enrichment <1e-10 and the log2 fold change averaged across all genes in the same pathway being > 0.1 by comparing 27 months to 6 months.

### Transcription Factor Motif Enrichment

The motif enrichment was performed with HOMER (for Hypergeometric Optimization of Motif EnRichment) software 4.10 (Heinz et al., 2010), which identifies motifs that are statistically enriched in the promotor region of a given list of genes. We applied the HOMER software using motifs of 6-12bp length over a range of 1000bp upstream and 50bp downstream of each transcriptional start site, using the lists of up-regulated or down-regulated genes for each tissue. Age-regulated genes were defined as described above.

## Data availability

Raw sequencing reads are available at the NIH Sequence Read Archive under the BioProject accession number **PRJNA516151**.

## ACKNOWLEDGMENTS

This work was supported by Novartis. We thank Judith Knehr, Ulrike Neumann, Marc Altofer, Kayhan Akyel, Lin Fan, Yanqun Wang and Oleg Iartchouk for performing RNAseq. Thank you to Bret Morin and Audrey Gray for collecting and processing tissues. All authors were employees of Novartis when this work was conducted, and some are current stockholders of Novartis (e.g., D.J.G). We thank the entire Age-Related Disorders group, the Chemical Biology & Therapeutics group, and the NIBR community for their enthusiastic support.

## AUTHOR CONTRIBUTIONS

TS and DJG conceived the study and designed experiments; TS, MM, JF, SW, WZ, HT and GR performed experiments and/or generated data. TS, MM, JF and RMG analyzed data. TS, MM, RMG and DJG wrote the paper. All authors approved the submission.

## Supplementary Figure Legends

**SUPPLEMENTARY FIGURE 1.**
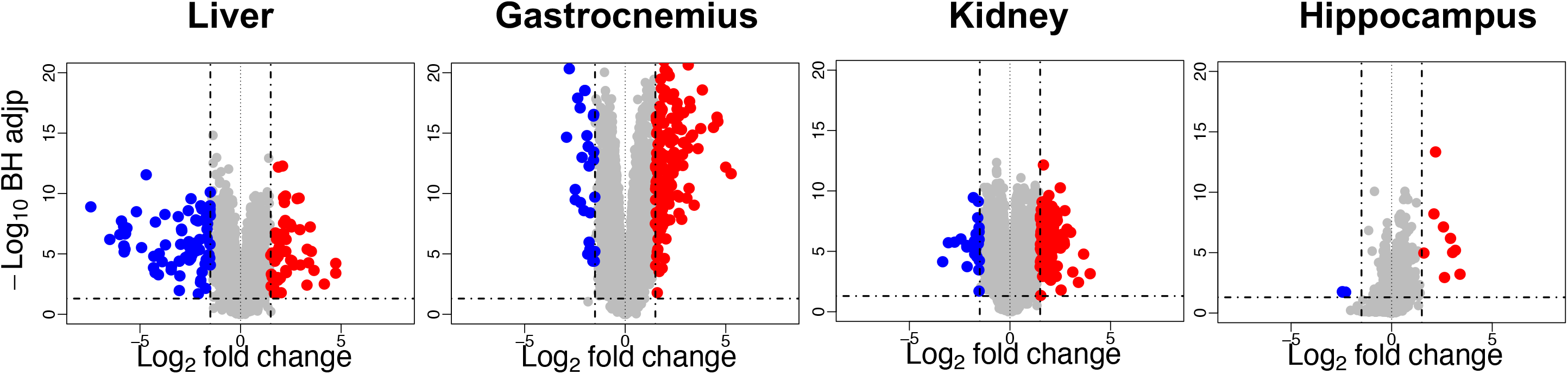
Volcano plots showing changes in gene expression in liver, gastrocnemius muscle, kidney and hippocampus between 27 month old and 6 month old rats (n=7-9 rats per group). Genes are represented by circles. X-axis represents log2 of fold change old versus young. Y-axis represents −log10 of the adjusted p value. Dotted lines indicate a Benjamini-Hochberg adjusted p-value of 0.05 (horizontal lines) and a fold change of 1.5 (vertical lines).

**SUPPLEMENTARY FIGURE 2.**
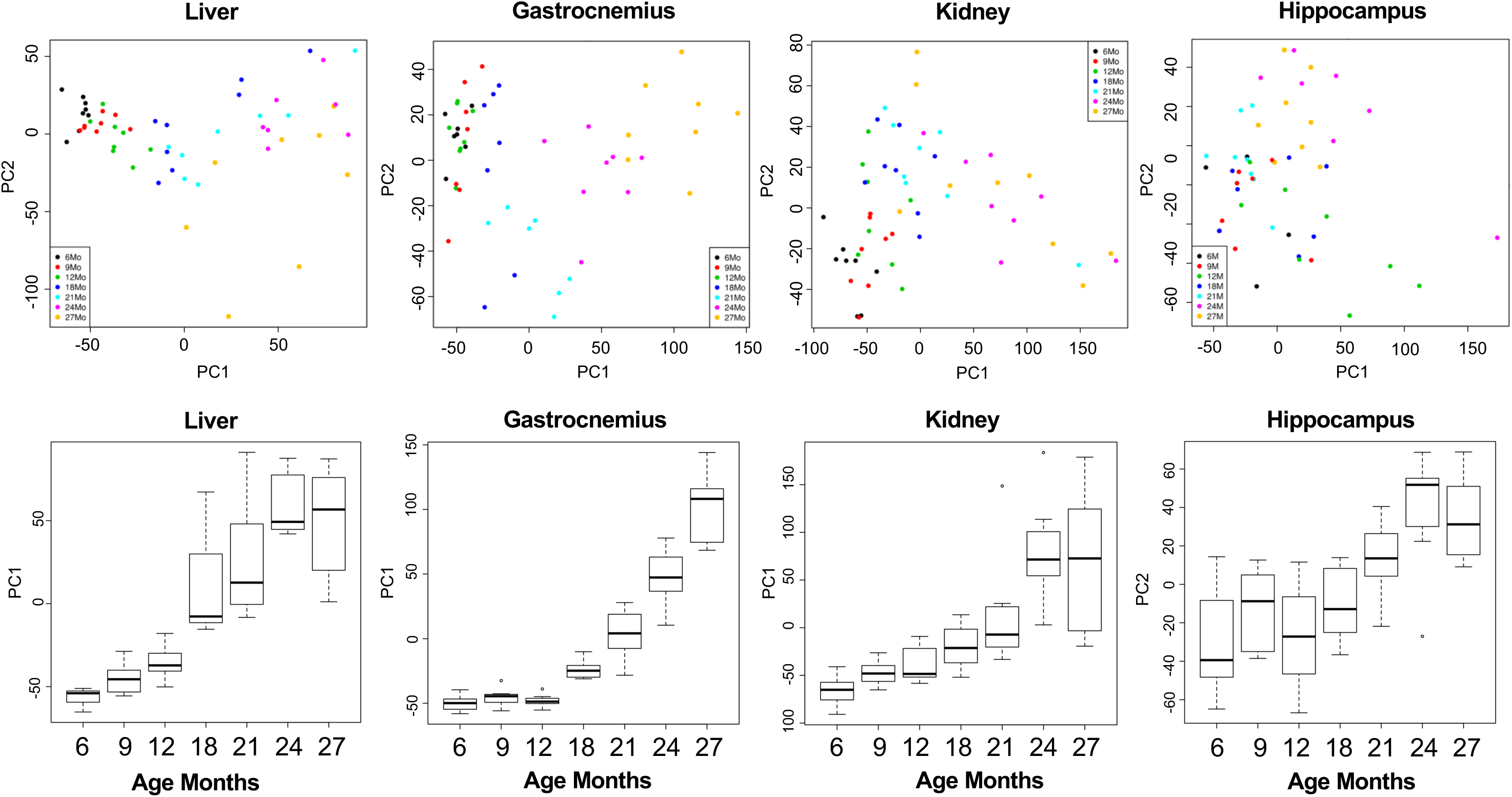
Principal component analyses for liver, gastrocnemius muscle, kidney and hippocampus RNAseq datasets (n= 7-9 rats per group). In a bottom panel, PCs that best correlated with age are shown: for liver, gastrocnemius muscle and kidney this was PC1 and for hippocampus PC2.

**SUPPLEMENTARY FIGURE 3.**
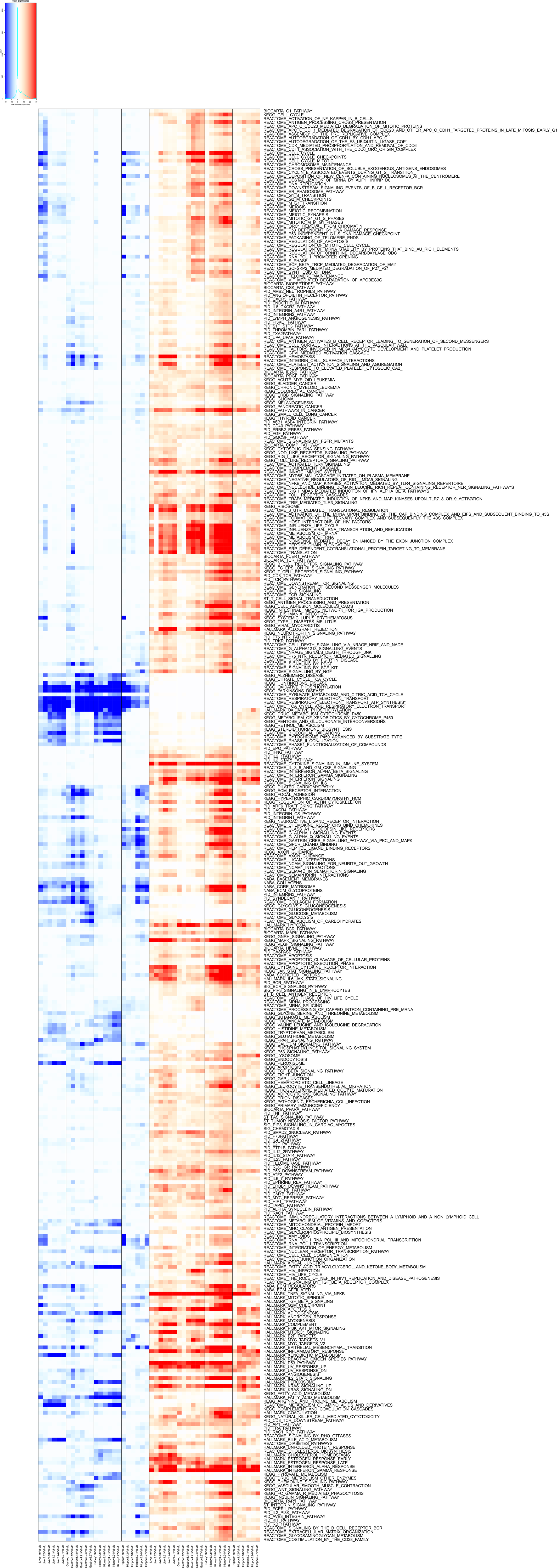
Identification of pathways UP- or DOWN- regulated with aging in liver, gastrocnemius muscle, kidney and hippocampus with gene set enrichment analysis. Pathway analyses are based on n=7-9 rats per group. Each row represents a pathway, annotated on the right hand side and each column represents a specific age comparison (i.e. 9 months vs 6 months, 12 months vs 6 months, etc up to 27 months vs 6 months). Colors represent statistical significance of enrichment of the pathways. Color transition from white to red shows up-regulation, while color transition from white to blue shows down-regulation of the pathway, with the color key indicating statistical significance of log10 of the p-value of enrichment.

**SUPPLEMENTARY FIGURE 4.**
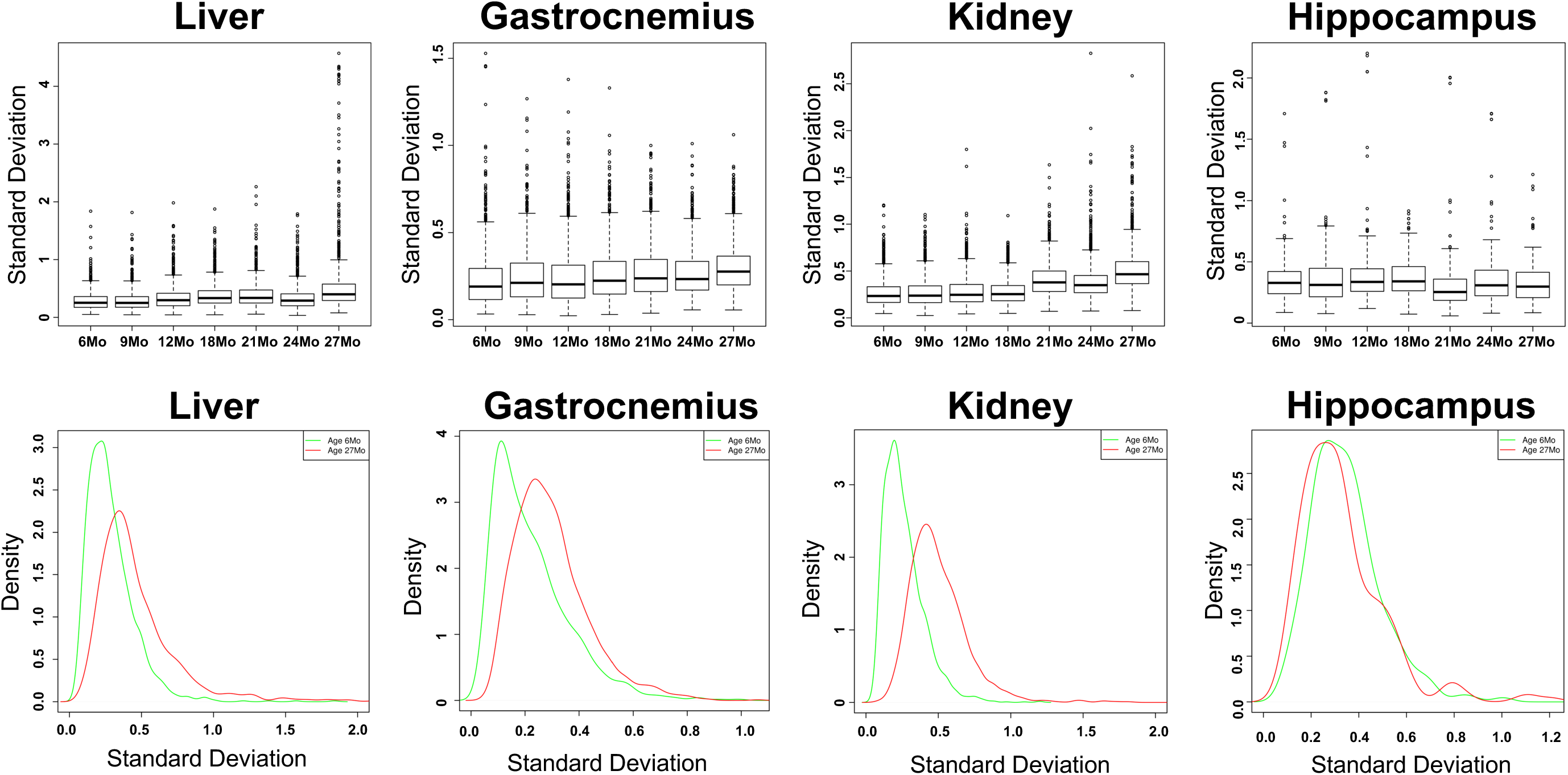
Gene expression variability with increasing age in liver, gastrocnemius muscle, kidney and hippocampus (n=7-9 rats per group). For each tissue, the standard deviation of age-regulated genes is plotted as a function of time as a boxplot (upper panel). The lower panel shows line density plots for standard deviation distributions of age-regulated genes in 6 month and 27 month old rats.

