## Supplemental Figures and Legends for "Age-related Gene Expression Signatures (AGES) in rats demonstrate early, late, and linear transcriptional changes from multiple tissues"

#### Supplementary Figure Legends

SUPPLEMENTARY FIGURE 1. Volcano plots showing changes in gene expression in liver, gastrocnemius muscle, kidney and hippocampus between 27 month old and 6 month old rats (n=7-9 rats per group). Genes are represented by circles. X-axis represents log2 of fold change old versus young. Y-axis represents -log10 of the adjusted p value. Dotted lines indicate a Benjamini-Hochberg adjusted p-value of 0.05 (horizontal lines) and a fold change of 1.5 (vertical lines).

SUPPLEMENTARY FIGURE 2. Principal component analyses for liver, gastrocnemius muscle, kidney and hippocampus RNAseq datasets (n= 7-9 rats per group). In a bottom panel, PCs that best correlated with age are shown: for liver, gastrocnemius muscle and kidney this was PC1 and for hippocampus PC2.

SUPPLEMENTARY FIGURE 3. Identification of pathways UP- or DOWN- regulated with aging in liver, gastrocnemius muscle, kidney and hippocampus with gene set enrichment analysis. Pathway analyses are based on n=7-9 rats per group. Each row represents a pathway, annotated on the right hand side and each column represents a specific age comparison (i.e. 9 months vs 6 months, 12 months vs 6 months, etc up to 27 months vs 6 months). Colors represent statistical significance of enrichment of the pathways. Color transition from white to red shows up-regulation, while color transition from white to blue shows down-regulation of the pathway, with the color key indicating statistical significance of log10 of the p-value of enrichment.

SUPPLEMENTARY FIGURE 4. Gene expression variability with increasing age in liver, gastrocnemius muscle, kidney and hippocampus (n=7-9 rats per group). For each tissue, the standard deviation of age-regulated genes is plotted as a function of time as a boxplot (upper panel). The lower panel shows line density plots for standard deviation distributions of age-regulated genes in 6 month and 27 month old rats.

### Supplementary Figure 1

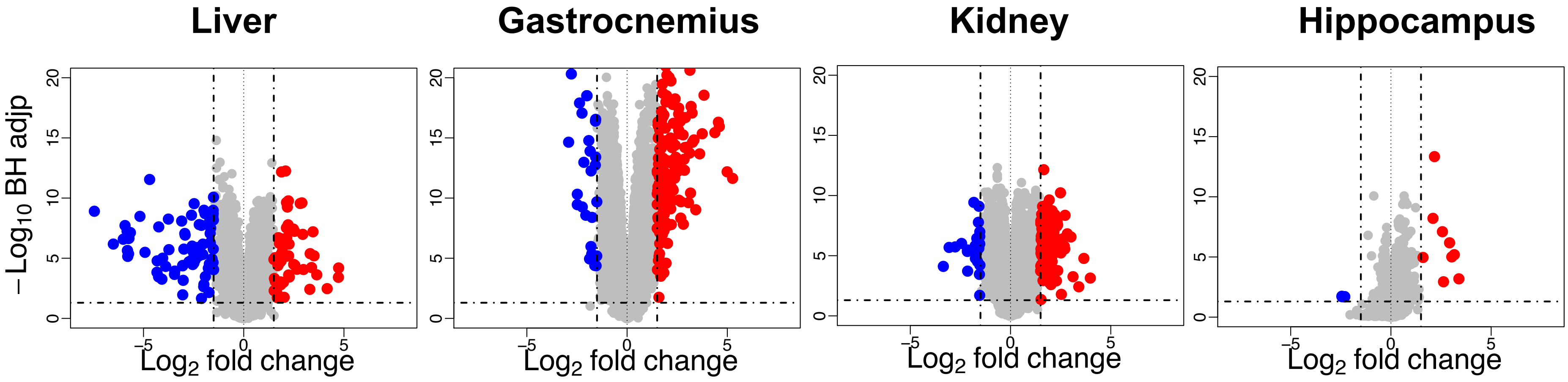

### Supplementary Figure 2

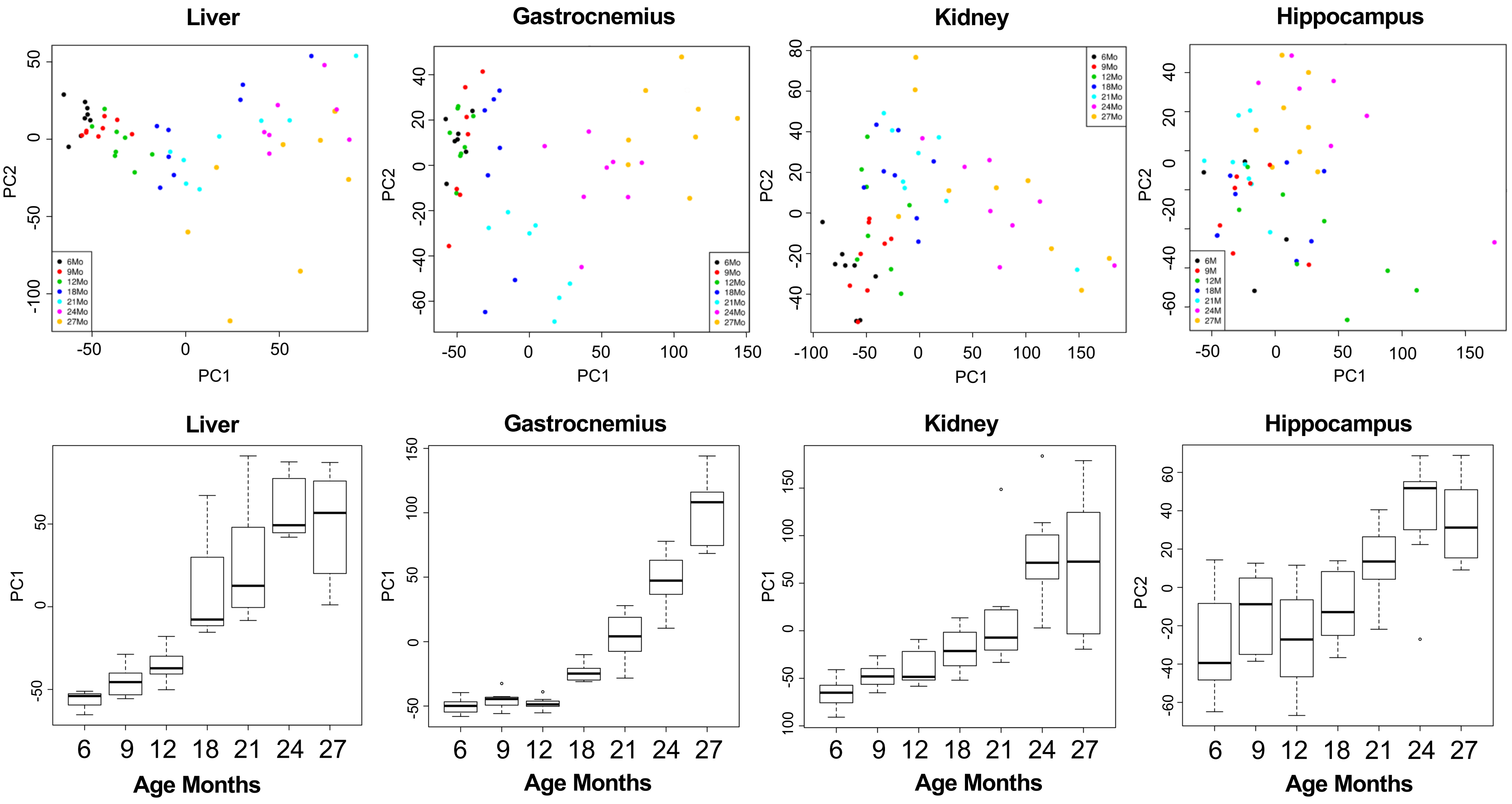

#### Supplementary Figure 3

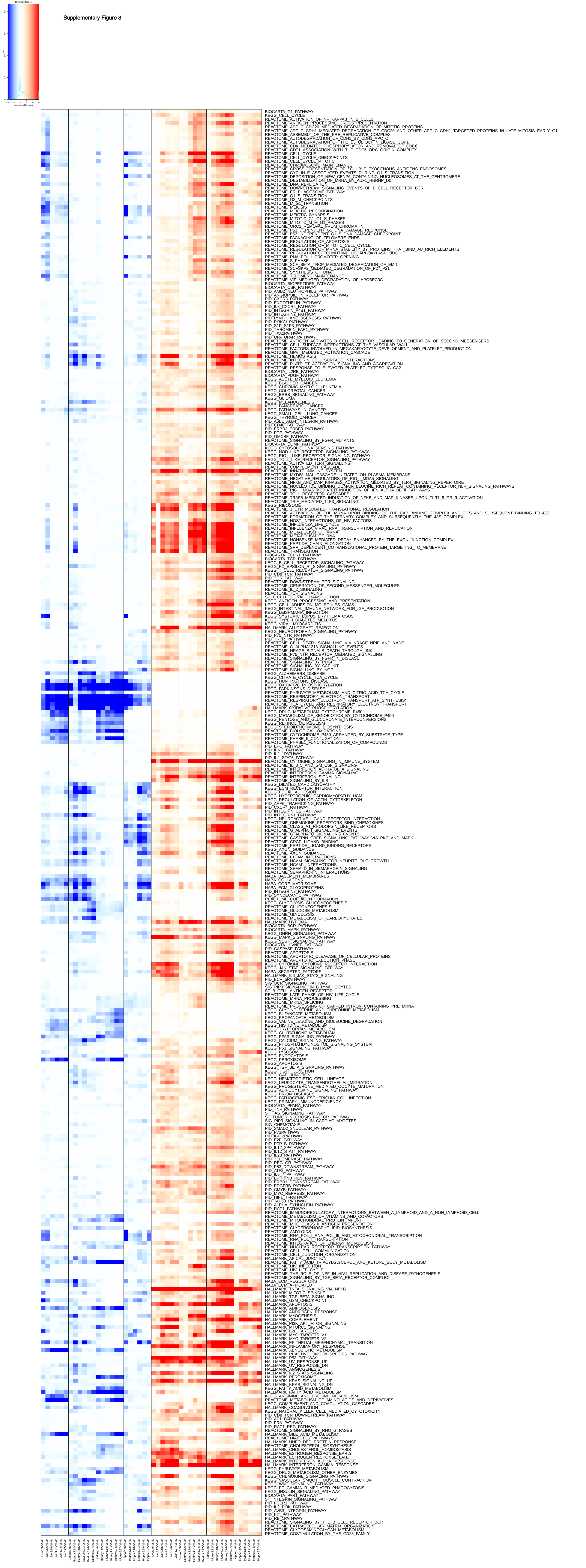

### Supplementary Figure 4

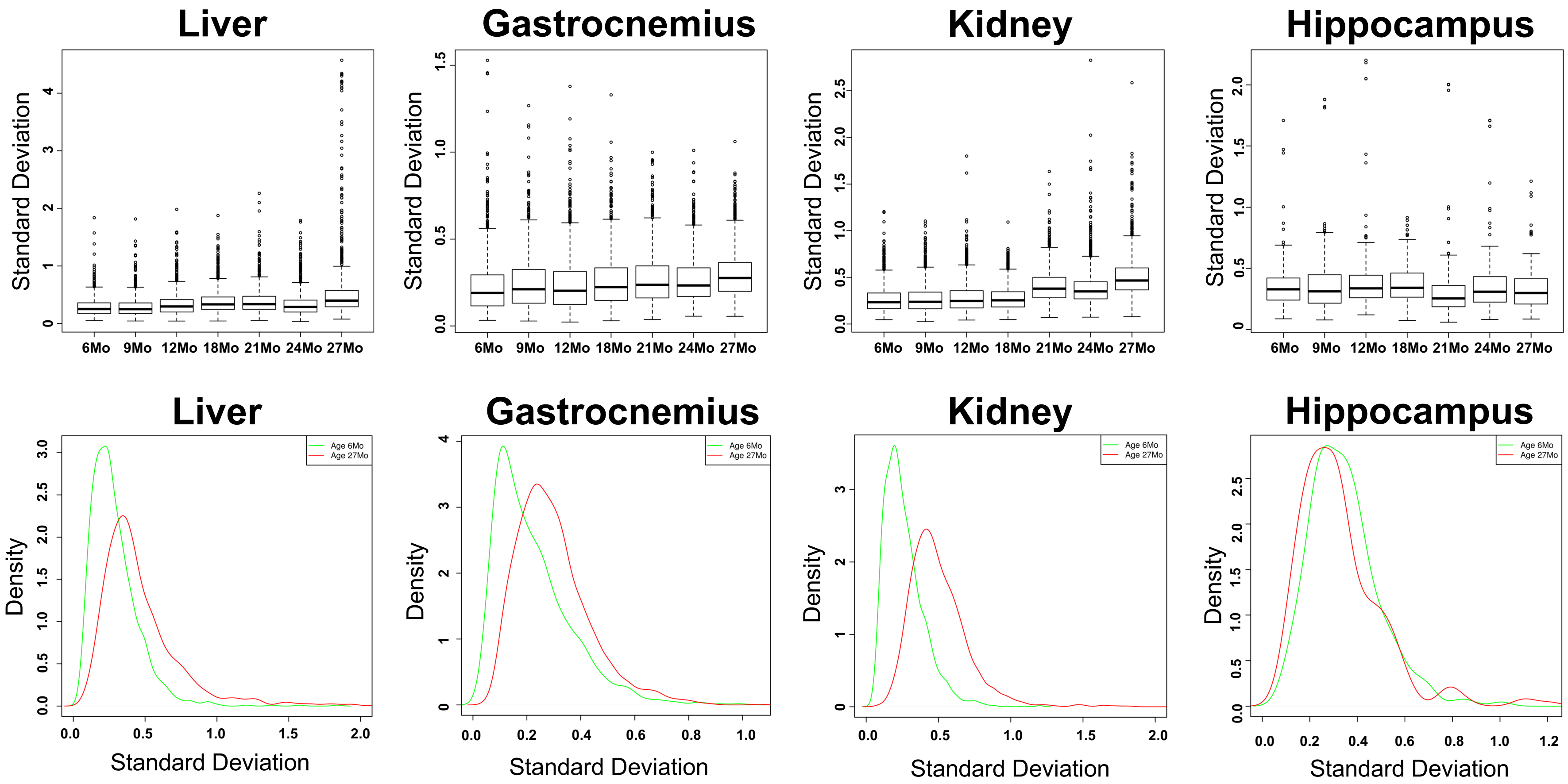
