## Supplementary Table 5 for "Age-related Gene Expression Signatures (AGES) in rats demonstrate early, late, and linear transcriptional changes from multiple tissues"

**Supplementary Table 5**. Summary of average reads (million) sequenced and mapped for each tissue.

|  | **Liver** | **Gastrocnemius** | **Kidney** | **Hippocampus** |
| --- | --- | --- | --- | --- |
| Average Reads Sequenced (million) | 44.6 | 36.6 | 48.5 | 52.0 |
| Average Reads Aligning to gene transcripts (million) | 26.1 | 18.5 | 22.6 | 18.9 |
